# Conformational states control Lck switching between free and confined diffusion modes in T cells

**DOI:** 10.1101/446732

**Authors:** Geva Hilzenrat, Elvis Pandžić, Zhengmin Yang, Daniel J. Nieves, Jesse Goyette, Jérémie Rossy, Katharina Gaus

## Abstract

T cell receptor (TCR) phosphorylation by Lck is an essential step in T cell activation. It is known the conformational states of Lck control enzymatic activity; however, the underlying principles of how Lck finds its substrate in the plasma membrane remain elusive. Here, single-particle tracking is paired with photoactivatable localization microscopy (sptPALM) to observe the diffusive modes of Lck in the plasma membrane. Individual Lck molecules switched between free and confined diffusion in resting and stimulated T cells. Conformational state, but not partitioning into membrane domains, caused Lck confinement as open conformation Lck was more confined than closed. Further confinement of kinase-dead versions of Lck suggests that Lck interacts with open active Lck to cause confinement, irrespectively of kinase activity. Our data supports a model that confined diffusion of open Lck results in high local phosphorylation rates and closed Lck diffuses freely to enable wide-range scanning of the plasma membrane.

---

T cell signaling is a tightly controlled process involving both simultaneous and sequential spatiotemporal events, involving membrane remodeling and redistribution of key signaling proteins’ ^1,2^. Engagement of the T cell receptor (TCR) with an antigenic pMHC on the surface of an antigen-presenting cell (APC) leads to the formation of immunological synapses and initiates downstream signaling events that lead to T cell activation ^4^ The Src family kinase Lck plays a crucial role in the signaling cascade. TCR engagement results in the membrane release ^5^ and phosphorylation of the immunoreceptor tyrosine-based motifs (ITAMs) located in the cytoplasmic tails of the CD3ζ chain by Lck ^6^. Phosphorylated sites on the TCR-CD3 complex become docking sites for the zeta chain-associated protein kinase 70 (ZAP70), that is further phosphorylated by Lck ^7^ before recruiting other proteins in the signaling cascade that are necessary for complete T cell activation.

The role of Lck in T cell activation as a signaling regulator is of particular interest due to its dynamic characteristics. Lck is a 56 kDa protein comprised of a Src homology (SH) 4 domain at the N-terminus, followed by a unique domain, an SH3 domain, an SH2 domain, a kinase domain and short C-terminal tail. Lck is anchored to the plasma membrane through its SH4 domain via post-translational acylation on three specific sites: a myristoylated Gly2 ^8^ and palmitoylated Cys3 and Cys5. The latter two are crucial for membrane binding and biological activity, enabling Lck diffusion in the inner leaflet of the plasma membrane and its recruitment to the immunological synapse ^9^ The unique domain interacts with the CD3ε subunit in the TCR-CD3 complex ^10^ as well as the co-receptors CD4 and CD8 ^11^ via zinc-mediated bonds. However, Lck does not require the co-receptors for recruitment to the immunological synapse or for TCR triggering ^12^, suggesting that freely diffusing Lck is sufficient for T cell activation.

Lck conformation is regulated by the phosphorylation of two tyrosine residues: Tyr^394^, where phosphorylation increases Lck activity, and Tyr^505^, whose phosphorylation reduces Lck activity ^13,14^ Intramolecular interactions between the phosphorylated Tyr^505^ (pTyr^505^) and the SH3 and SH2 domains cause rearrangements that keep Lck in a closed, inactive conformation ^15,16^. When dephosphorylated by CD45, Lck exists in an open, primed conformation. When Tyr^394^ is trans-autophosphorylated ^14^, rearrangements in the activation loop stabilize the active conformation ^17^. Lck’s diffusion behavior ^18^ and conformational state ^19,20^ are thought to be regulated by the activation state of the cell. The conformational state also influences Lck clustering ^21^. This means that not only does Lck conformational state regulate Lck enzymatic activity but also aids in its diffusive search strategy.

Whether Lck becomes ‘active’, i.e., converted into the open conformation upon TCR engagement, has been controversial. There is evidence of global changes in relative populations of closed and open Lck in resting *versus* stimulated T cells ^19,20^. These studies propose that Lck undergoes conformational changes upon T cell activation, driving it from its closed state to an open state, therefore enhancing its activity. Using biochemical analyses, conformational heterogeneity was observed in resting and stimulated T cells ^22^, suggesting a “standby-model” in which ~40% of Lck is in the open conformation in both resting and stimulated T cells. Ballek et al. challenged these observations in a later report that used different cell lysis conditions ^13^. Another study, based on measurements of fluorescence resonance energy transfer (FRET) between fluorescent proteins fused to the N- and C-terminals of Lck, concluded there was no significant change in open *versus* closed populations of Lck even after T cell stimulation ^24^. While different papers report different percentages of open Lck in pre-stimulated cells, constitutively active Lck were also found in CD8^+^ memory T cells and may account for the enhanced sensitivity to antigen in these cells ^25^. A pool of active Lck existing prior to T cell stimulation led to the idea that rapid TCR triggering post receptor engagement may be caused by changes in Lck spatial rearrangements as opposed to, or in addition to conformational changes. Using single-molecule localization microscopy in fixed cells, we previously showed that Lck distributed differently on the cell surface depending on its conformational state ^21^, with open Lck residing preferentially in clusters and closed Lck preventing clustering, regardless of the cell’s activation state. However, this study only captured the overall distribution of open or closed Lck and the movement of Lck clusters, but to understand the search strategy of the membrane-bound kinase the dynamic behavior of individual molecules need to be taken into account.

The efficiency of a dual-state search strategies has previously been demonstrated in other systems ^26^. For Lck, such a strategies would entail that individual molecules oscillate between two distinct states: a confined state that corresponds to high Lck activity and a diffusive state that enables the kinase to scan the membrane for substrates. Such a dual-state search strategy may account for the high fidelity of Lck-mediated phosphorylation of the available TCR-CD3 complexes while also retaining high signaling sensitivity when membrane-detached cytosolic tails of the CD3 complex are limited. The former would be mediated by the high enzymatic activity in Lck clusters while the high level of diffusion of Lck in the closed state would enable the latter.

The dynamic behavior of Lck was previously mapped with single particle tracking (SPT) in live cells, revealing, for example, the differences in Lck diffusion in stimulated *versus* resting T cells and the formation of microclusters, but without linking dynamics to conformational states ^18,27^. Overall changes in diffusion constants were observed, as well as segregation into different confinement zones, attributed to actin and other proteins compartmentalizing the membrane ^27,28^ or to the formation of membrane microdomains ^18^. Recently, Lck compartmentalization upon TCR stimulation was attributed to the formation of close-contact zones between the T cell membrane and the stimulating surface, possibly because of exclusion of CD45, in line with the kinetic segregation model ^29^. These works, however, did not take into account the conformational change in Lck.

In the current study, we utilize SPT using photoactivatable localization microscopy (sptPALM)^30^ as a tool to study the diffusion of wild type (WT) and mutated Lck, lacking the tyrosine residues on positions 394 and 505, to measure the dynamics of the closed and open forms, respectively ^19,20^. Lck variants were tagged with photoactivatable monomeric cherry (PAmCherry) ^31^, expressed in Jurkats 1.6E cells and imaged in resting and activating conditions. Single trajectories were extracted and analyzed in order to find periods when the proteins underwent confined diffusion and the fraction of confined versus free proteins was determined ^32^. Measurements of different Lck variants showed that conformation has a key role in Lck’s substrate search strategy, with the open form dwelling more in confinements compared to the closed form. Further, we provide evidence of Lck-Lck interaction in the open conformation in stimulated T cells. Taken together, the data suggest that Lck continuously switched between open and closed states, which is likely to determine the probability of productive encounters between Lck and its substrates.

## Results

### Identification of free and confined states of Lck in live T cells

In order to characterize the diffusion patterns of Lck, we applied single particle tracking on image sequences from different experimental conditions and decomposed each trajectory into free and confined segments. Jurkat E6.1 cells were transfected with either wild-type Lck (wtLck) fused to PAmCherry (wtLck-PAmCherry) or a truncated construct of Lck containing only the first ten amino acids that are responsible for Lck anchoring to the membrane (Lck10-PAmCherry). T cells were stimulated for ~5 minutes at 37°C on a coverslip coated with anti-CD3 and anti-CD28 antibodies and imaged either in live-cell conditions or after chemical fixation. In each experiment we acquired 10,000 frames with an 18 ms exposure for the duration of ~197 s. Imaging was done while continuously photo-activating and exciting the fluorophores. Trajectories shorter than 15 frames and immobile particles (particles with a low RMSD, as described in the Methods section) were excluded from analysis.

We decomposed trajectories by adopting a previously described post-tracking analysis ^32^. Briefly, every trajectory is first fragmented into overlapping windows. For each window, the normalized variance of the location of the particle is calculated as a measure of the level of confinement, L_Conf_, according to:

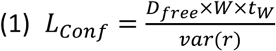

where D_free_ is the diffusion coefficient of freely diffusing Lck in μm^2^ sec^-1^, W is the window size in frames, tW is the temporal length of the window in seconds and var(r) is the variance in μm^2^. We chose the diffusion coefficient of Lck10-PAmCherry as the value for D_free_ for all versions of Lck as Lck10 is membrane anchored but does not interact with other proteins. We then defined a threshold of L_Conf_ above which particles are considered confined. For this threshold, we chose the most abundant L_Conf_ value for wtLck-PAmCherry in stimulated T cells, following the procedure published previously ^32^ (dotted line in Fig. 1A). This threshold was suitable because the majority of values for Lck10-PAmCherry in resting cells were below this threshold and all values for wtLck-PAmCherry in fixed cells were above the threshold (Fig. 1A). In order to ensure that Lck molecules were in fact confined, rather than just temporarily slowing down, we only regarded a molecule as confined if it has an L_Conf_ value above the threshold for three or more consecutive windows. Trajectories were then segmented into confined and free periods (Fig. 1B), depending on whether L_Conf_ was above or below the threshold (Fig. 1C). From this analysis it was evident that molecules that diffused slowly for 3 or more consecutive states were found to be confined (Fig. S1). This analysis was applied to all trajectories recorded in a cell (Fig. 1D). As is evident from this sptPALM analysis, individual wtLck-PAmCherry molecules in live cells switched between free and confined diffusive states (Fig. 1D) while in fixed cells, only confined or immobile molecules were observed (Fig. 1D).

**Fig.1.**
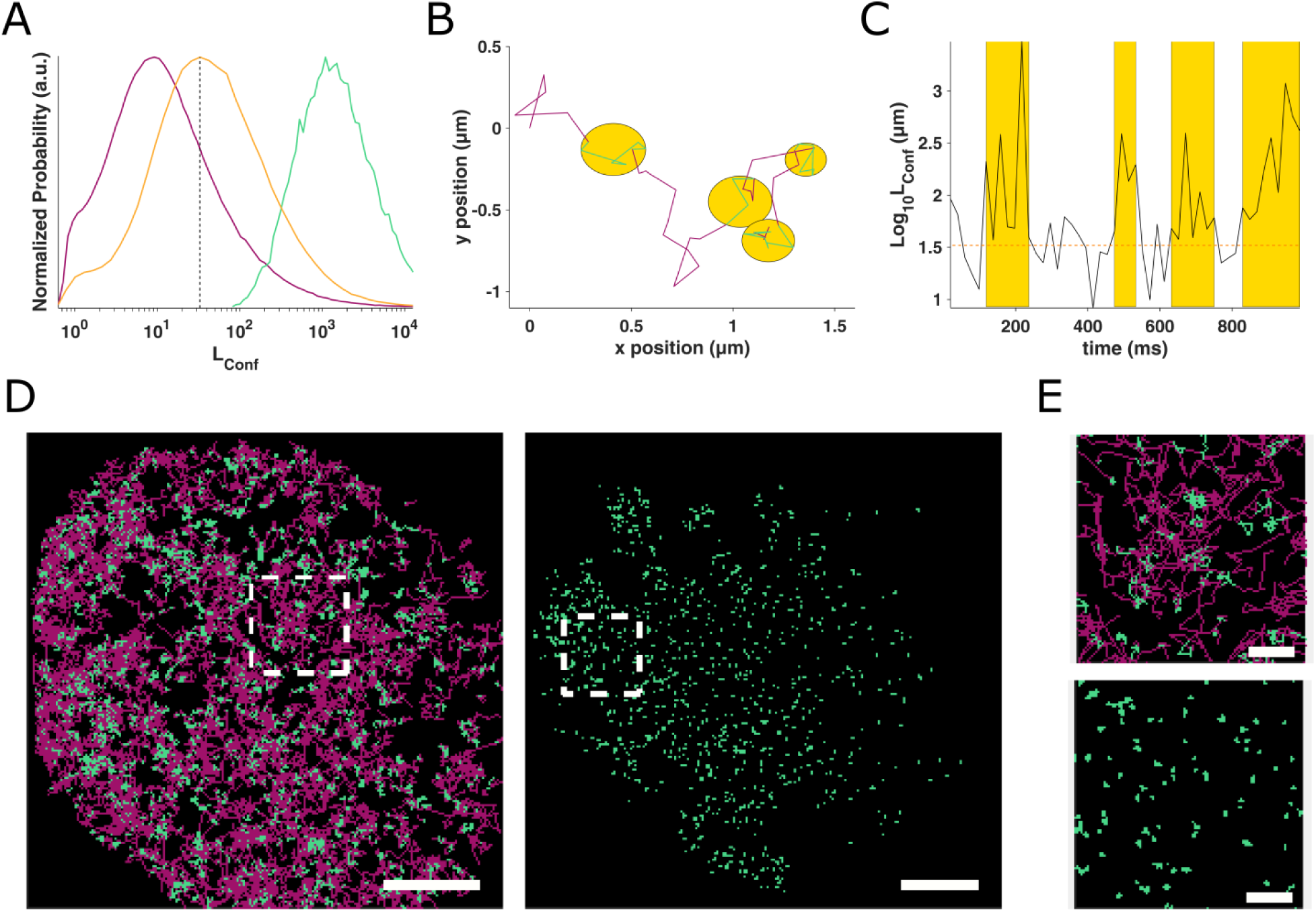
wtLck switches between free and confined states. **(A)** L_conf_ acquired for Lck10-PAmCherry in resting Jurkat cells (purple), wtLck-PAmCherry in stimulated Jurkat cells (orange) and wtLck-PAmCherry in fixed cells (cyan), normalized to peak value. The dashed vertical line marks the threshold where a particle was to be considered confined, i.e., if it had three or more consecutive steps with an L_Conf_ value greater than that threshold. **(B)** An experimental trajectory decomposed to free (magenta) and confined (cyan) states, with the confinements highlighted in yellow circles. **(C)** Time evolution of L_Conf_ values for the trajectory in (B) with the threshold marked with an orange dashed line and the confined periods with a yellow shade. **(D)** Trajectory decomposition maps of wtLck-PAmCherry in a stimulated live cells (left) and fixed Jurkat cells (right) Free periods are colored magenta, whereas confined periods are colored cyan. Scale bar = 5 μm. **(E)** 5 μm by 5 μm zoomed-in regions of interest in (D) (top – live, bottom - fixed). Scale bar = 1 μm.

### Wild-type Lck was more confined in stimulated than resting T cells

Previous studies provided evidence that T cell activation decreases the overall diffusion of Lck ^18,27^ In our experiments, resting T cell data was generated by placing T cells expressing wtLck-PAmCherry onto coverslips coated with anti-CD90 antibodies. This resulted in good T cell adhesion, but not TCR signaling or T cell activation ^33^. Our measurements confirmed the decrease in diffusion (Fig 2A; Movie S1: resting - right, stimulated - left), with diffusion coefficients of 1.16 μm^2^ s^-1^ (1.15-1.17) to 0.69 μm^2^ s^-1^ (0.68-0.7) for resting and stimulated cells, respectively (Fig. 2A, Fig. S2a; Movie S1). We wanted to assess whether this slowdown is caused by enhanced spatial compartmentalization in the membrane. Thus, we conducted the L_Conf_ analysis for wtLck-PAmCherry in resting and stimulated cells. When comparing the L_Con_f histogram of wtLck in stimulated T cells (Fig. 2B, blue) *versus* resting T cells (Fig. 2B, orange), it is noticeable that the values in activated cells are shifted to higher values, resulting in a mean L_Con_f value of 32.9 in stimulated cells and 29.1 in resting cells.

**Fig.2.**
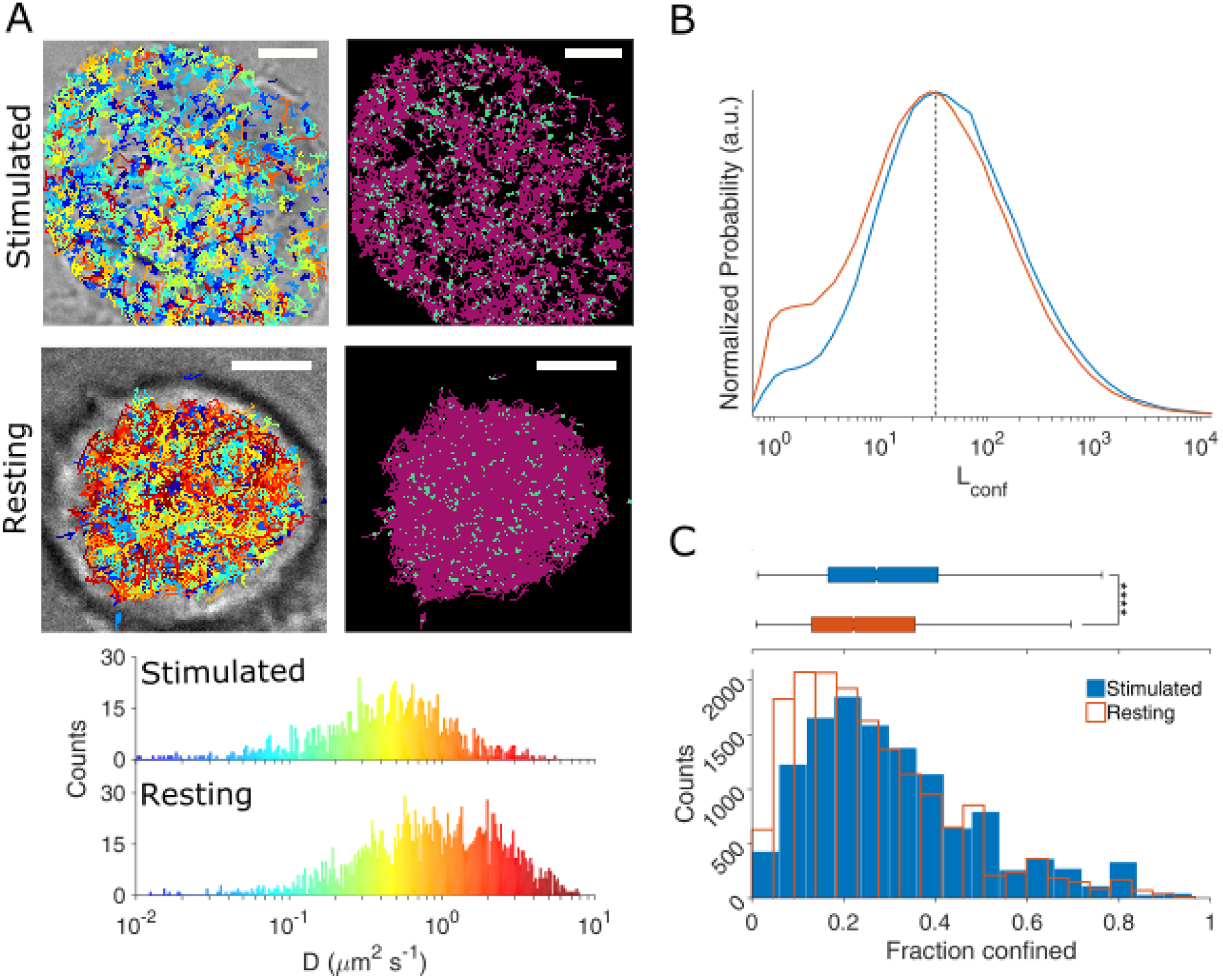
wtLck-PAmCherry is more confined in stimulated cells. (A) Representative stimulated and resting Jurkat E6-1 cells expressing wtLck-PAmCherry. The left panels show bright field images of the cells with detected trajectories overlaid, color-coded according to their initial diffusion. The right panels show the free (magenta) and confined (cyan) modes of diffusion. Scale bar = 5 μm. Bottom: diffusions histogram corresponding to the cells above, sharing mutual color-coding. **(B)** L_Conf_ histograms for wtLck-PAmCherry in resting (orange) and stimulated (blue) cells. **(C)** Histograms of the fraction of confined wtLck-PAmCherry molecules obtained for 13 stimulated (blue) and 17 resting (orange) Jurkat cells. Box plot shows the median. Notch 95% confidence interval, box edges first and third quartile, lines Tukey’s fences, **** p≤0.00001.

Next we examined whether the decrease in local displacement variance is due to a redistribution of wtLck-PAmCherry into confinements that would result in an increase in the number of consecutive steps that fall above the L_Con_f threshold value. Thus, we segmented the total video into segments of five frames (Fig. S3), in which we asked how many particles, out of the total number of particles imaged, were confined. Histograms obtained for stimulated and non-stimulated cells (Fig. 2C) were collected. There was a clear difference in peak value for the two populations as well as a larger tail of high values for wtLck-PAmCherry in stimulated cells. As a consequence, the populations were statistically different (Fig. 2C) when tested against the null hypothesis according to which the samples are drawn from the same population, using the rank sum test, with different medians and non-overlapping 95% confidence intervals with the values of 27.27% (26.67-27.78) and 22.22% (21.82-22.73) for stimulated and resting cells, respectively. The percentage of confined wtLck-PAmCherry were 30.97% (30.63-31.3) and 26.4% (26.14-26.68) in stimulated and resting cells, respectively.

Overall, these results show that wtLck-PAmCherry diffused slower in stimulated cells compared to resting cells, suggesting that in addition to a global reduction in diffusion, a redistribution of Lck into confinements had occurred. These results are in agreement with an increase in wtLck-PAmCherry clustering in fixed stimulated *versus* fixed resting T cells ^21^.

### Membrane anchoring alone is not contributing to Lck confinement

Lck confinement may be attributed to the formation of membrane domains, i.e., changes in membrane order, as a result of TCR triggering ^34^ If that is the case, a truncated version of Lck, Lck10, that includes the first ten amino acids that are responsible to Lck anchoring to the membrane as it contains the post-translational lipid modifications, is expected to experience the same slowdown and confinement as full-length Lck. However, we did not observe such a scenario (Fig. 3A; Movie S2), as the diffusion coefficients found for Lck10-PAmCherry in stimulating and resting conditions remained high (Fig. S2b). The overall level of confinement of Lck10-PAmCherry was almost identical for both resting and stimulated cells, with a peak L_Conf_ value of 7.2 and 7.74, respectively (Fig. 3B). These values were significantly different from the ones found for wtLck-PAmCherry, with most of the probability function having a value below the threshold described above. A histogram of confinement events (Fig. 3C) shows comparable peak values for both stimulated and resting cells. No statistically significant difference was found between the two samples (Fig. 3C, top panel), as shown by median of 9.62% (9.43-9.8) and 9.68% (9.52-10.00) for Lck10-PAmCherry expressed in stimulated and resting cells, respectively. Further, the mean fraction of confined particles was slightly higher in resting cells, with values of 13.96% (13.74-14.18) and 14.73% (14.45-15.00) for stimulated and resting cells, respectively. These values were lower than those found for wtLck-PAmCherry, suggesting Lck10-PAmCherry was far less confined than wtLck-PAmCherry, even in stimulated cells.

**Fig. 3.**
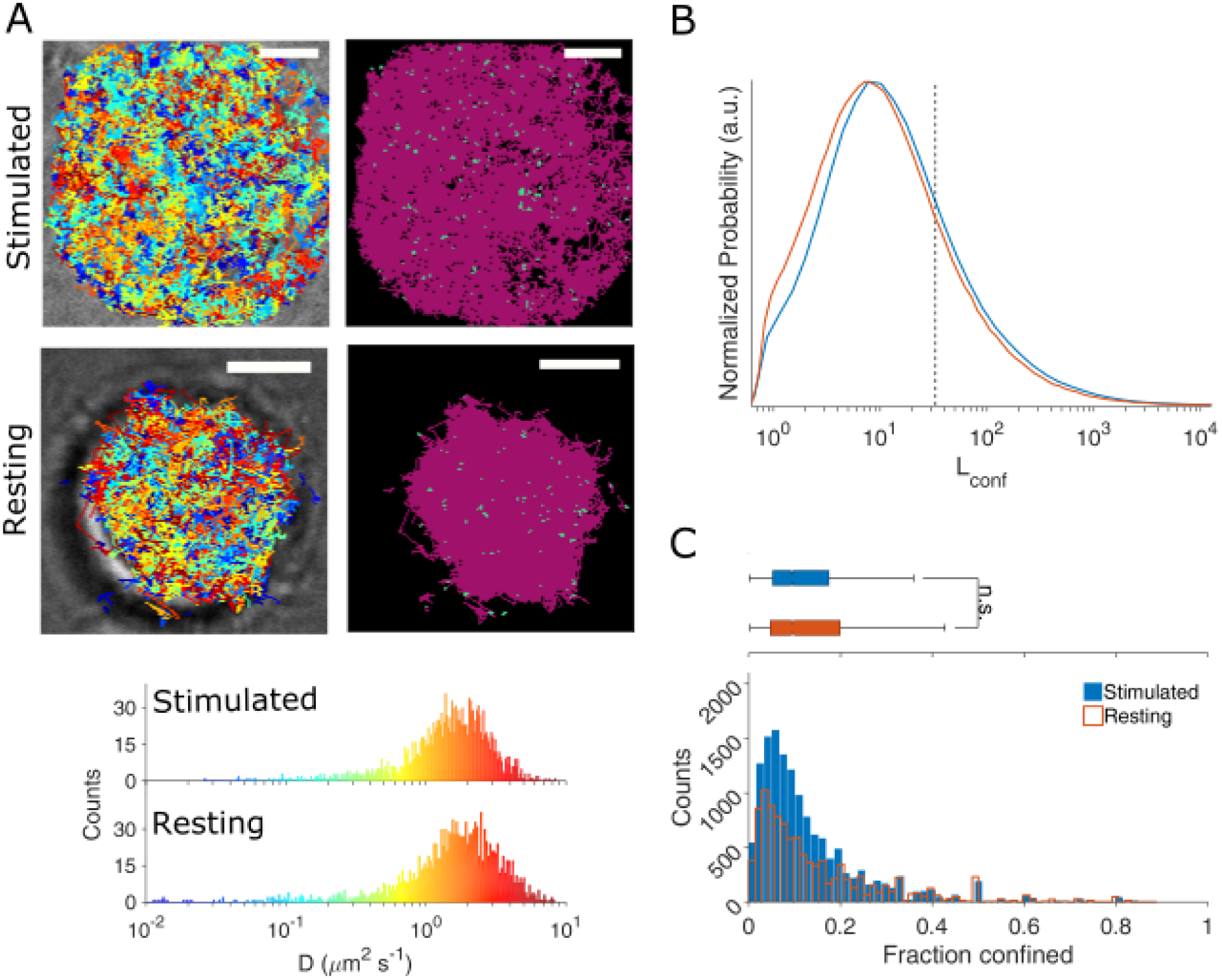
Lck10-PAmCherry demonstrates free-diffusion in resting and stimulated calls. **(A)** Representative stimulated and resting Jurkat E6-1 cells expressing Lck10-PAmCherry. The left panels show bright field images of the cells with detected trajectories overlaid, color-coded according to their initial diffusion. The right panels show the free (magenta) and confined (cyan) modes of diffusion. Scale bar = 5 μm. Bottom: diffusions histogram corresponding to the cells above, sharing mutual color-coding. **(B)** L_Conf_ histograms for Lck10-PAmCherry in resting (orange) and stimulated (blue) cells. **(C)** Histograms of the fraction of confined Lck10-PAmCherry molecules obtained for 19 stimulated (blue) and 15 resting (orange) Jurkat cells. Box plot shows the median. Notch 95% confidence interval, box edges first and third quartile, lines Tukey’s fences, n.s. p>0.01

Taken together, the data strongly suggest that the increased confinement observed for full-length wtLck-PAmCherry was not due to global changes in membrane organization or membrane domains ^18^ as confinement of Lck10 in resting and stimulated T cells was similar.

### Open Lck is highly confined in stimulated and resting cells

Previously, we reported that Lck clustering was regulated by the kinase’s conformational state ^21^. We thus quantified the influence of conformation on confinement of Lck in live cells. First, we introduced a Tyrosine-to-Phenylalanine mutation at position 505 in Lck (Lck^Y505F^). The mutation prevents the binding of Lck pTyr^505^ to its own SH2 domain. This mutation is well known as ‘constitutively open’ ^19-21,24,35^ and ‘hyperactive’ ^13^. An overall change in the diffusion constants due to cell activation (Fig. 4A; Movie S3) was observed, with values of 0.65 μm^2^ s^-1^ (0.64-0.66) and 0.95 μm^2^ s^-1^ (0.94-0.96) in stimulated and resting cells, respectively (Fig. S2c). Further, L_Conf_ values for Lck^Y505F^-PAmCherry were higher than that of wtLck-PAmCherry (Fig. 4B), with peak values of 39.28 and 42.53 in stimulated and resting cells, respectively, with <50% of log_10_(L_Conf_) events above the confinement threshold. In contrast to wtLck-PAmCherry, the L_Conf_ distributions of Lck^Y505^-PAmCherry were similar in resting and stimulated T cells. This similarity was also observed in the histograms of the confined fractions (Fig. 4C), with a large population of Lck^Y505^ molecules falling into the right tail of the distribution. Importantly, unlike in the corresponding data for wtLck-PAmCherry, these values were not significantly different from each other (Fig. 4C, top), with median values and overlapping 95% confidence interval of 26.55% (26.32-26.67) and 26.39% (26.14-26.67) for stimulated and resting cells, respectively. The means of Lck^Y505^ were 29.85% (29.59-30.11) and 29.97% (29.74-30.22) in stimulated and resting cells, respectively.

**Fig. 4.**
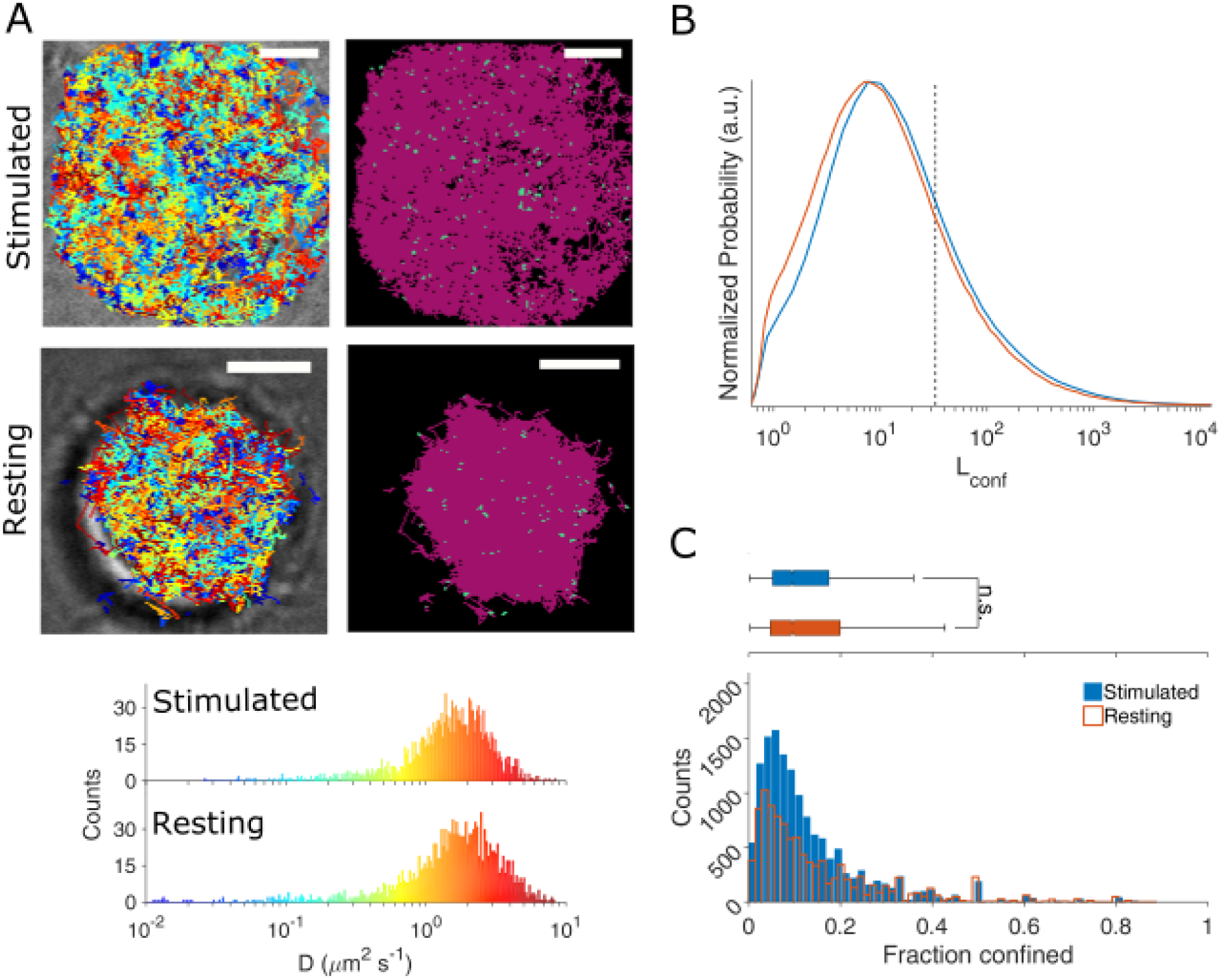
Lck^Y505F^-PAmCherry is equally confined in stimulated and resting cells. (A)Representative stimulated and resting Jurkat E6-1 cells expressing Lck^Y505F^-PAmCherry. The left panels show bright field images of the cells with detected trajectories overlaid, color-coded according to their initial diffusion. The right panels show the free (magenta) and confined (cyan) modes of diffusion. Scale bar = 5 μm. Bottom: diffusions histogram corresponding to the cells above, sharing mutual color-coding. (B) L_Conf_ histograms for Lck^Y505F^-PAmCherry in resting (orange) and stimulated (blue) cells. (C) Histograms of the fraction of confined Lck^Y505F^-PAmCherry molecules obtained for 14 stimulated (blue) and 18 resting (orange) Jurkat cells. Box plot shows the median. Notch 95% confidence interval, box edges first and third quartile, lines Tukey’s fences, n.s. p>0.01.

These data show that when Lck is locked in the open state, it is also driven into a more confined diffusive behavior, which is comparable with wtLck-PAmCherry in stimulated cells (Fig. S4). Although the diffusion coefficient found for Lck^Y505F^-PAmCherry is lower, in terms of confinement, open Lck was insensitive to the T cell activation with Lck^Y505F^-PAmCherry confinement levels being similar in both stimulated and resting cells. This indicates that Lck confinement is driven by the open conformation of the kinase and supports that a higher proportion of Lck is in the open conformation in stimulated T cells ^19,20,36^.

### Closed Lck is as confined as wild-type Lck in resting cells

To further investigate the hypothesis that Lck conformation regulates Lck diffusive behavior, we expressed a closed form of Lck in Jurkat cells. A mutation in position 394 converting a tyrosine into phenylalanine (Lck^Y394F^) prevents the formation of the activation loop and results in reduced-activity ^14^ or an inactive Lck ^13^ because of the hyper-phosphorylated tyr^50522^ that constitutively closes the enzyme ^19^.

As with the wtLck and Lck^Y505F^, Lck^Y394F^-PAmCherry did undergo a decrease in diffusion coefficient due to stimulation (Fig.5A; Movie S4), from 1.24 μm^2^ s^-1^ (1.22-1.26) in resting cells to 0.88 μm^2^ s^-1^ (0.87-0.89) in stimulated cells (Fig. S2d). We applied the same sptPALM analysis to Lck^Y394F^-PAmCherry and lower L_Conf_ values were obtained with peak values of 32.93 and 30.36 in stimulated and resting cells, respectively (Fig. 5B). Histograms of the fraction of confined Lck^Y394F^-PAmCherry showed the populations were skewed towards lower values (Fig. 5C). Similarly to Lck^Y505F^-PAmCherry, Lck^Y394F^-PAmCherry showed no statistically significant difference between stimulated and resting cells (Fig. 5C, top panel) and medians of 22.22% (21.88-22.58) and 21.95% (21.43-22.22) for Lck^Y394F^-PAmCherry in stimulated and resting cells, respectively. The mean confinement fractions were 26.09% (25.85-26.33) and 26.24% (25.92-26.55) for Lck^Y394F^-PAmCherry in stimulated and resting cells, respectively.

**Fig. 5.**
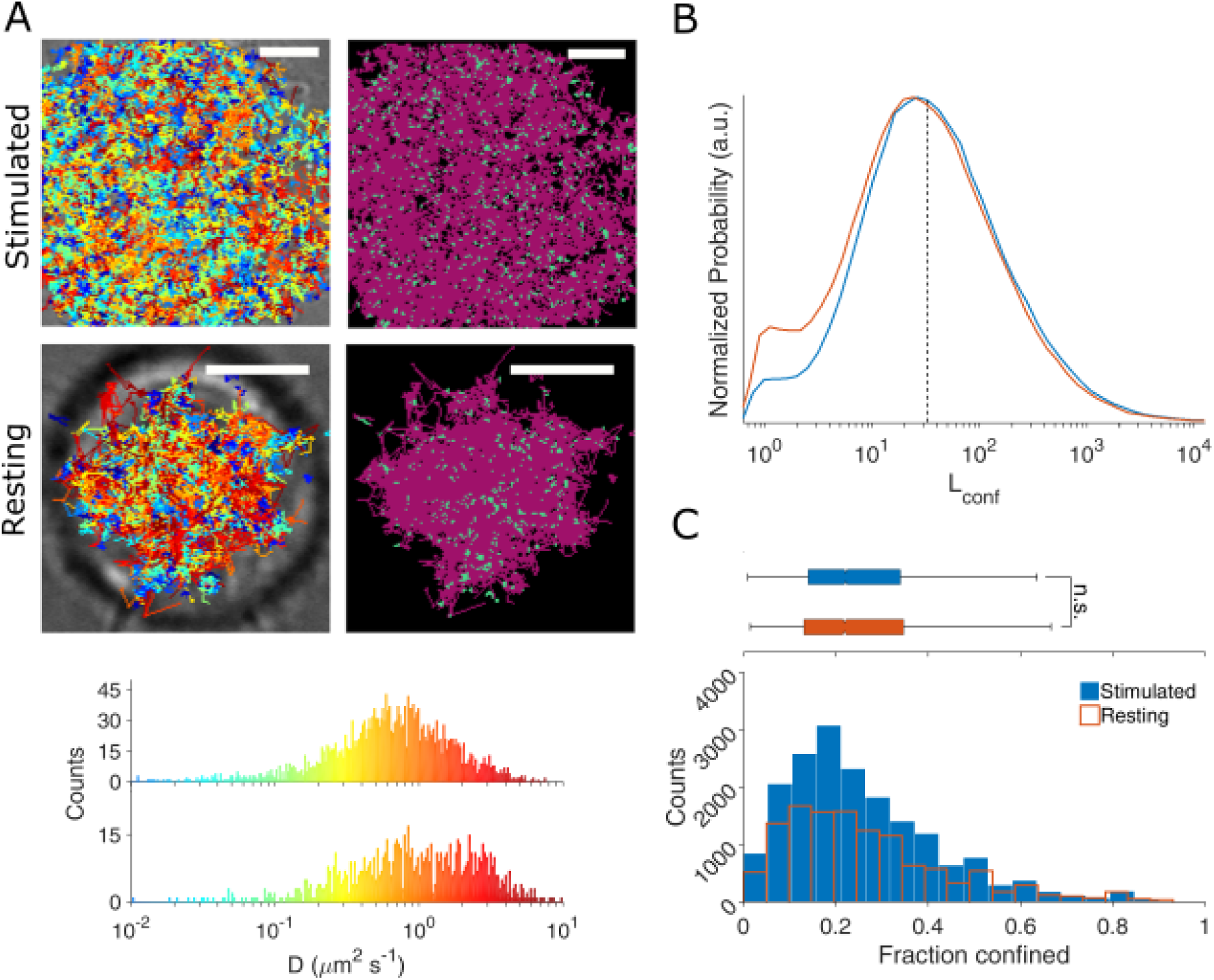
Lck^Y394F^-PAmCherry is equally confined in stimulated and resting cells. **(A)**Representative stimulated and resting Jurkat E6-1 cells expressing Lck^Y394F^-PAmCherry. The left panels show bright field images of the cells with detected trajectories overlaid, color-coded according to their initial diffusion. The right panels show the free (magenta) and confined (cyan) modes of diffusion. Scale bar = 5 μm. Bottom: diffusions histogram corresponding to the cells above, sharing mutual color-coding. **(B)** L_Conf_ histograms for Lck^Y394F^-PAmCherry in resting (orange) and stimulated (blue) cells. **(C)** Histograms of the fraction of confined Lck^Y394F^-PAmCherry molecules obtained for 16 stimulated (blue) and 14 resting (orange) Jurkat cells. Box plot shows the median. Notch 95% confidence interval, box edges first and third quartile, lines Tukey’s fences, n.s. p>0.01.

The confinement fraction values we found for the closed Lck were smaller than the ones found for the open Lck (Fig. S4), illustrating the significance conformational states have on Lck diffusion. Indeed closed Lck has a similar level of confinement as wtLck in resting cells while open Lck was similarly confined as wtLck in activated cells (Fig. S4). Thus, the data confirms that confinements are regulated by the conformational state of Lck with open Lck being more confined and closed Lck being less confined.

### Lck self-associates with other Lcks, depending on its conformation and activity

An open Lck that is also phosphorylated in Tyrosine 394 is known to be active, while studies done on Lck^Y505F^ showed hyperactivity ^13,14^ By expressing a constitutively inactive Lck, we could assess whether Lck confinement relies on enzymatic activity. We tested an Lck variant in which the lysine in position 273 in the kinase domain is replaced with Arginine (Lck^K273R^-PAmCherry, Fig. 6, Fig. S5), which has been shown to render Lck kinase-dead ^37^. Different diffusion coefficients of 0.82 μm^2^ s^-1^ (0.81-0.83) and 1.13 μm^2^ s^-1^ (1.12-1.15) were observed for Lck^K273R^-PAmCherry in stimulated and resting cells, respectively (Fig. S2e). However, similar L_Con_f histograms, with values of 34.80 for stimulated and 37.58 for resting cells, were obtained (Fig. 6B, blue and orange) with no significant difference observed in the fraction of time spent confined (Fig 6C, blue and orange). Additionally, Lck^K273R^-PAmCherry spent 25.80% (25.56-26.07) and 25.62% (25.38-25.86) of time confined in stimulated and resting cells, respectively (Fig 6C). Thus, the level of confinement kinase-dead Lck did not depend on the T cell activation status as it did for wtLck (Fig. S5). Assuming that the K273R mutation disables Lck activation via autophosphorylation, as hypothesized previously ^13^, the finding suggest that confinement of wtLck in stimulated T cells is regulated by Lck activation.

**Fig. 6.**
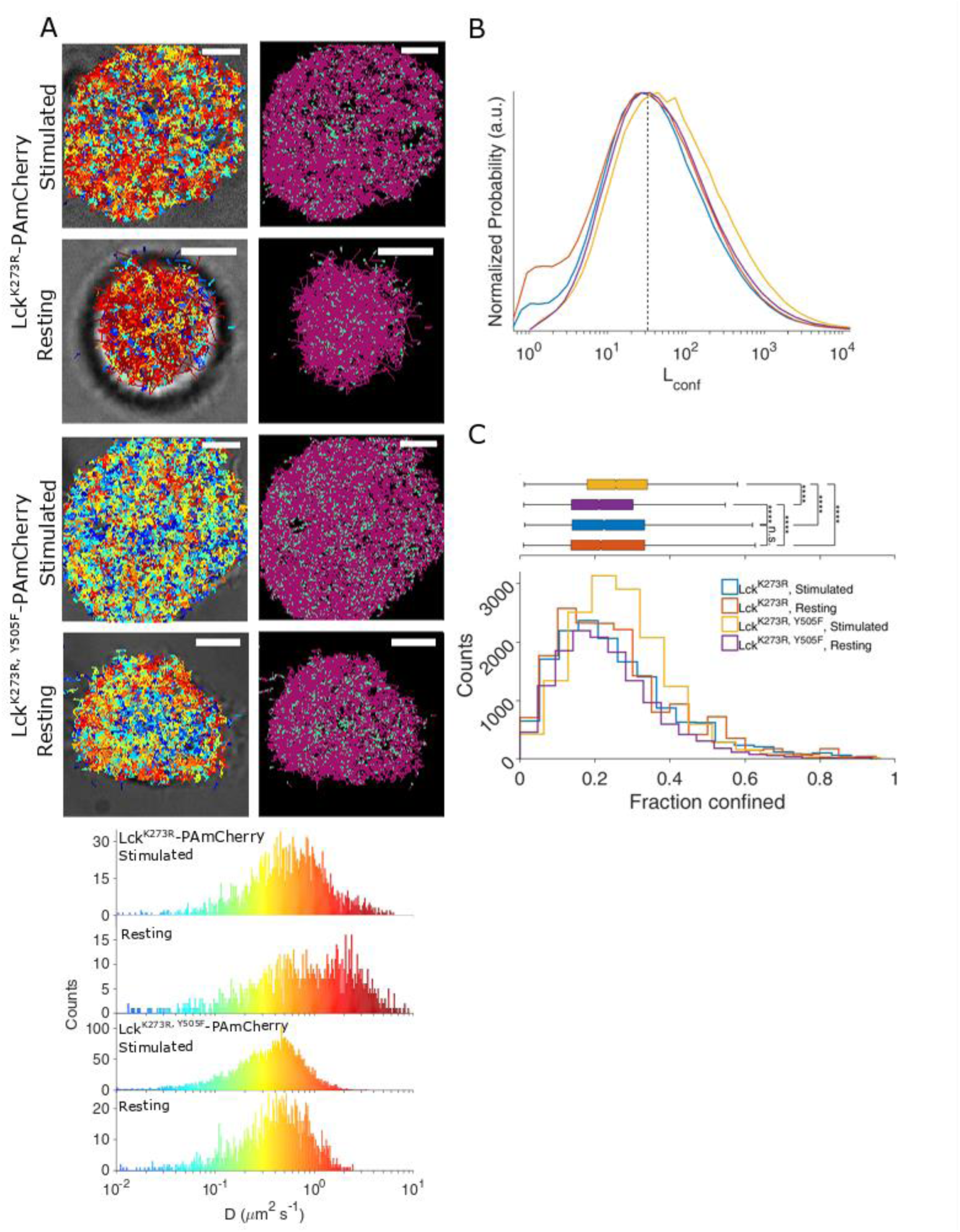
Confinement analyses for Lck^K273R^-PAmCherry and Lck^K273R^’ ^Y505F^-PAmCherry in stimulated and resting cells. **(A)** Representative stimulated and resting Jurkat E6-1 cells expressing Lck^K273R^-PAmCherry and Lck^K273R^’ ^Y505F^-PAmCherry. The left panels show bright field images of the cells with detected trajectories overlaid, color-coded according to their initial diffusion. The right panels show the free (magenta) and confined (cyan) modes of diffusion. Scale bar = 5 μm. Bottom: diffusions histogram corresponding to the cells above, sharing mutual color-coding. **(B)** L_Conf_ histograms for Lck^K273R^-PAmCherry and Lck^K273R,Y505F^-PAmCherry in and stimulated cells. (orange, blue, purple and yellow, respectively). **(C)** Histograms of the fraction of confined Lck^K273R^-PAmCherry molecules obtained for 12 stimulated (blue) and 14 resting (orange) Jurkat cells and histograms of the fraction of confined Lck^K273R,Y505F^-PAmCherry obtained for 8 stimulated (yellow) and 8 resting (purple) Jurkat cells. Box plot shows the median. Notch 95% confidence interval, box edges first and third quartile, lines Tukey’s fences, **** p≤0.00001, n.s. p>0.01.

To further test this hypothesis, we expressed a constitutively-open kinase-dead mutant Lck^K273R^’ ^Y505F^-PAmCherry. This mutant had slower diffusion coefficients of 0.41 μm^2^ s^-1^(0.41-0.42) and 0.51 μm^2^ s^-1^ (0.5-0.51) in stimulated and resting cells, respectively (Fig. 6A; Fig. S2f; Movie S5), values that were slower than those obtained for Lck^K273R^-PAmCherry (Fig. S2e, f). Further, Lck^K273R^’ ^Y505F^-PAmCherry had higher L_conf_ values in stimulated cells (Fig. 6C, purple and yellow) compared to resting cells (44.78 and 35.09, respectively).

Conducting the same analysis to quantify confinement fractions, we found a large fraction of kinase-dead mutant in the open conformation was highly confined in stimulated cells (Fig. 6C, purple and yellow). When comparing total trajectories, Lck ^K273R, Y505F^-PAmCherry in stimulated cells was more confined than in resting cells and more than Lck^K273R^ in both cell activation statuses (Fig. S5). These data confirm the conclusion that open, but not necessarily enzymatically active Lck confined the kinase in distinct zones in the plasma membrane.

Lck^K273R, Y505F^-PAmCherry was more confined in stimulated cells (26.97% (26.76-27.18)) than resting cells (23.30% (23.08-23.52)). It is possible that the open, kinase-dead variant of Lck interacts with endogenous Lck in Jurkat cells that were already shown to be confined in stimulated cells (Fig. 2). This would suggest that open Lck is confined in activated T cells by Lck-Lck interaction. Moreover, the lowered confinement for the K273R-Y505F mutant in resting cells compared to stimulated cells excludes the possibility of confinement due to increase in hydrodynamic radius of the enzyme (Fig. S5). Taken together, the experiments with the kinase-dead version of Lck confirmed the finding that it was the open conformation that caused the Lck confinement. Thus it is likely that the enzyme switches between open and close conformation, which results in a dual-state search strategy where open and active Lck is confined, and closed and inactive Lck diffuses freely (Fig. 7).

**Fig. 7.**
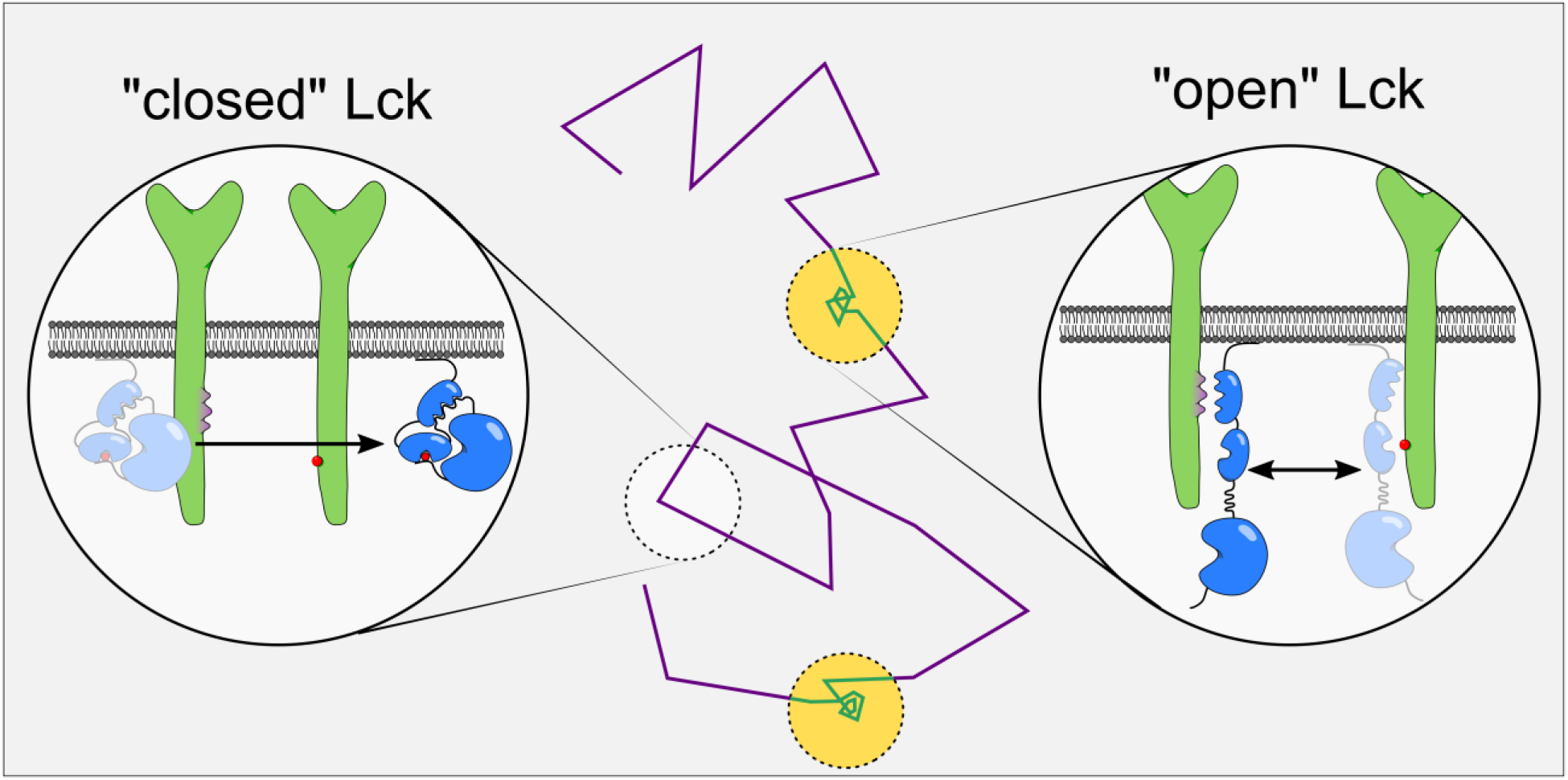
Two-stage diffusion model of Lck that combines an efficient search strategy with a high phosphorylation rates of substrates. Lck (illustrated in blue) exists in two main conformations: a closed conformation characterized by low catalytic activity and mediated by intramolecular interactions; and an open conformation characterized by high catalytic activity and free SH2 and SH3 domains. Our data propose that the closed conformation diffuses unimpeded (purple line), whereas the open conformation interacts with other membrane proteins (illustrated in green) via SH2 and SH3 domain mediated interactions and becomes confined (yellow circles) through rapid rebinding (teal line). Free diffusion allows Lck to scan large membrane areas while confinement in the open conformation enables high substrate phosphorylation rates.

## Discussion

Phosphorylation of the TCR-CD3 complex by the kinase Lck is an essential step in T cell activation ^38^. While it is relatively well documented that the conformational states control enzymatic activity, how membrane-bound Lck finds and phosphorylates its substrates is not well understood. For example, the link between phosphorylation state and activity is well established ^39^, as well as some interactions of Lck with other proteins ^40-43^ and lipids ^44^ Most studies so far have focused on whether or not T cell stimulation results in an ‘activation’ of Lck itself, i.e., whether there is an overall increase of Lck molecules in the open conformation and whether a stable pool of open Lck already exists in resting T cells. However, it is also possible that in the dynamic environment of the inner leaflet of the plasma membrane, Lck switches between open and closed states, as many other types of enzymes do ^45-47^. Utilizing single molecule localization microscopy (SMLM) techniques, our group showed that open Lck clusters were bigger and denser than closed Lck clusters ^21^. In SMLM, re-excitation of the same molecule can lead to overestimation of clustering ^48^. Thus, we investigated here whether Lck switches between confined and free diffusion modes. By tracking single Lck molecules, we were indeed able to set a threshold to distinguish a population that diffuses freely from one that exhibited restricted diffusion. We found that wild-type Lck (wtLck-PAmCherry) transitioned between free and confined states in both resting and stimulated T cells, strongly suggesting that the kinase has a sophisticated search strategy.

In a study employing immunofluorescence staining, a pre-existing pool of constitutively active Lck was used to explain the readiness of Lck to phosphorylate the TCR immediately after T cell stimulation ^22^, while showing no difference in the fraction of open Lck when comparing stimulated to resting cells. Therefore, it was speculated that Lck undergoes re-distribution upon T-cell stimulation, while maintaining the same overall fraction of Lck in the open and closed conformation. By examining the diffusion modes of Lck, as a function of conformational status, we can provide an alternative explanation of how the kinase can be efficient at both searching for substrates and phosphorylating the TCR complex. Firstly, we found that T cell stimulation significantly changed the behavior of wtLck, promoting Lck molecules to spend more time in confinements, compared to resting cells. Further our results showed that in resting cells, wtLck behaved like the closed Lck mutant in both activating and resting conditions. In contrast, in stimulated cells, wtLck demonstrated a diffusion pattern that was similar to that of the open Lck mutant in both conditions. These observations led us to the conclusion that T cell activation leads to a higher proportion of open Lck, supporting the recent findings that were obtained with a fluorescence resonance energy transfer (FRET) Lck biosensor ^19^ Our findings do not exclude the possibility of a pre-existing pool of open Lck.

Comparing the level of confinement of open and closed Lck mutations (Fig. S4) clearly showed that diffusion behaviour dependent on the conformational state of the enzyme. Lck^Y394F^-PAmCherry i.e. closed Lck was confined than wtLck-PAmCherry in stimulated cells and Lck^Y505F^-PAmCherry i.e. open Lck in stimulated and resting cells. Further, Lck^Y394F^-PAmCherry demonstrated similar confinement to that of wtLck-PAmCherry in resting cells. The values obtained for the open mutant, both in stimulated and resting cells were closer to the value that we obtained for wtLck-PAmCherry in stimulating conditions. Taken together, our data support that notion that the open conformational state of Lck is responsible for Lck confinement and that T cell activation resulted in converting some of the wtLck molecules into the open state ^19,20^.

All Lck variants demonstrated some level of confinement in resting conditions. As these results were obtained by expressing Lck variants in Jurkat cell lines, this confinement may be an outcome of self-association with endogenous active Lck, and may be related to a pre-existing pool of opened Lck in resting cells. Other mechanisms such as Lck’s SH4 domain interaction with lipid rafts ^49,50^ and microdomains ^51,52^ were previously suggested. Such scenarios should have, however, also affected Lck10-PAmCherry, as this segment is responsible for anchoring Lck to the membrane and should have resulted in slower diffusion in stimulated cells. However this was not the case; similarly to closed Lck (Lck^Y394F^-PAmCherry) we found no difference in confinement of Lck10-PAmCherry in resting and stimulated T cells. Further, one may hypothesize that Lck confinement is indirectly related to membrane domains, by interacting with other proteins that are lipid raft-associated. However, from our results with open Lck (Lck^Y505F^-PAmCherry) we could not find support for this idea, as Lck^Y505F^ was similarly confined in resting and stimulated cells.

The kinase-dead mutant, Lck^y^-PAmCherry, was found to be minimally-confined in resting and stimulated cells. It is possible that the K273R mutation in Lck prevents the rearrangements in the activation loop that prevent self-association of Lck^K273R^, or interaction with other proteins, thus, limiting confinement ^13^. Relying on our results obtained for wtLck-PAmCherry, and thus assuming that a greater population of endogenous Lck was in the open, confined state in stimulated Jurkat cells compared to resting cells, Lck^K273R, Y505F^-PAmCherry was found to be highly confined in stimulated cells, supporting the hypothesis that Lck self-associated with other active Lck, therefore, promoting a more confined population. This is consistent with a previous report on Lck self-association in the open conformation ^43^. Given that Lck in the open conformation exhibited confined diffusion and hyperactivity ^13,14^, it is highly likely that this state results in high local phosphorylation rates.

In conclusion, we provide evidence that the conformation of Lck was the main driver of Lck diffusion modes with open Lck causing confined diffusion and closed Lck enabling free diffusion. Individual Lck molecules can switch between confined and free diffusion in resting and stimulated T cells. This is consistent with a dual-state search strategy that enables Lck to scan large areas of the membrane in the closed state, but efficiently phosphorylate TCR-CD3 complexes at numerous sites in the open state.

## Methods and Materials

### Plasmids

Lck and Lck10 were amplified by PCR and inserted within the Ecot1 and Age1 restriction sites of a pPAmCherry-N1 plasmid. Y394F, Y505F and K273R mutations were further introduced via site-directed mutagenesis.

### Sample Preparation

Jurkat cells were cultured in RPMI medium (Gibco) containing phenol-red and supplemented with 10% (vol/vol) FBS, 2 mM L-glutamine (Invitrogen), 1 mM penicillin (Invitrogen) and 1 mM streptomycin (Invitrogen). Cell cultures were passaged normally every ~48 hours, when the cell count reached ~8×10^5^ viable cells per ml. The cells were cultured for at least 1 week (3-4 passages) after thawing prior to transfection and imaging. No cells were used after passage 20.

Cells were transfected by electroporation (Neon; Invitrogen); briefly, cells were collected before reaching a cell density of 8×10^5^ cell/ml and while ≥90% viable. The cells were washed twice with 1x PBS in 37°C and resuspended in the resuspension buffer (R-buffer) provided with the Neon kit. Three pulses of 1325 V with 10 ms duration were applied. The cells were allowed to recover in clear RPMI 1640 medium (Gibco) supplemented with 20% HI-FBS for overnight. Prior to imaging, fresh warm (37°C) media with 40 mM HEPES, pH 7.4 was added to achieve a final concentration of 20 mM HEPES.

1.5H coverslips (Marienfeld-Superior) were waterbath-sonicated in four 30-minutes stages: 1 M KOH, Acetone, EtOH and ultra-pure (18 MΩ) water. The coverslips were then allowed to adsorb 0.01% PLL (Sigma) in ultra-pure water for 15 minutes. Excess solution was later aspirated and the coverslips were baked-dry in 60 °C for 1 hour. Finally, after cooling-down, the coverslips were coated with either 0.01 mg/ml anti-CD3 (OKT3; eBioscience) and 0.01 mg/ml anti-CD28 (CD28.2; Invitrogen) for stimulating conditions or 0.01 mg/ml αCD90 for (Thy-1; eBioscience) for resting conditions and let rest in 4°C overnight before imaging. The coverslips were washed 3 times with phosphate buffer saline (PBS) pre-warmed to 37°C before the cells were transferred onto them to interact with the antibodies. For live-cell experiments, imaging took place ~5 minutes after cell-transfer, or fixed with 4% paraformaldehyde (P6148; Sigma) in 37°C, followed by 3 washing cycles with PBS for fixed-cell imaging.

### Imaging

For each sptPALM experiment 10,000 frames were acquired in a ~50 frames per second (18 ms exposure time) rate on a total internal reflection fluorescence (TIRF) microscope (ELYRA, Zeiss) in 37 °C using a 100× oil immersion objective (N.A. = 1.46) and a 67.5° incident beam angle. PAmCherry fused to Lck variants were continuously photoactivated using a 405 nm laser radiation tuned to 0.5-5 μW (interchangeable during acquisition to maintain a low density) and continuously excited with a 561 nm laser tuned to 2.5 mW. Point density was monitored by using ZEN (Zeiss) online-processing tool.

### Data Analysis

All accumulated data are comprised of three biologically-independent experiments, i.e. each mutant was imaged in two or more cells (in one of the three repetitions, where a repetition relates to a different transfection) in each cell-activation state (stimulated or resting). We used Diatrack ^53^ for fitting the point spread functions (PSFs) to a Gaussian with a 1.75 pixel width (1 pixel ≈ 0.097 nm) and then to track the particles by setting the search radius to 10 pixels. The data was later analyzed by a custom MATLAB (Mathworks) adaptation of the trajectory analysis part of a previously published multi-target tracing (MTT) code ^32^. Immobile particles (RMSD < 2 pixels) and trajectories shorter than 15 frames were excluded from analysis. Stages of confined and free diffusion were detected according to equation 1, with D_free_ = 2.15 μm^2^ /sec (Fig. S2b, bottom), W = 4 and t_W_ was the sum of the exposure time and the CCD reading time (~19.7 ms). To detect time spent in confinement, each sequence was segmented to non-overlapping windows of 5 frames and in each block of 5 frames, the ratio of confined:total particles was calculated. Each value of one 5 frames-window is a count in the histogram. All data processing and statistical analyses were performed in MATLAB.

### Statistical Tests

To compare between two populations of confinement fractions, that do not normally distribute, we used the Mann-Whitney U test, while the Kruskal-Wellis test was used for multiple datasets followed by a bonferroni post-hoc test. **** and n.s. indicate p≤0.00001 and p>0.01, respectively. Ranges around median and mean values in supplementary text are the 95% confidence intervals calculated from bootstrapping the data by sampling 10,000 times.

## Supplementary Materials

Fig.S1 Relationship between Lck diffusion coefficient and confinement

Fig.S2 Diffusion coefficients histograms of wtLck, Lck10, LckY505F, LckY394F, LckK273R and LckK273R, Y505F in stimulated and resting Jurkat cells

Fig.S3 Illustration of confinement ratio analysis

Fig.S4 Comparison of confinement analysis result

Fig.S5 Comparison of confinement analysis result of wtLck-PAmCherry, LckK273R-PAmCherry and LckK273R, Y505F-PAmCherry in stimulated and resting cells

Movie S1

Movie S2

Movie S3

Movie S4

Movie S5

## Author Contributions

GH performed experiments, modified analysis, analyzed data, and wrote manuscript. EP established analysis and helped write the manuscript. ZY was responsible for the generation of Lck constructs. DJN and JG aided in writing and drafting of the manuscript. JR provided guidance with experiments. KG designed the project, interpreted the data and wrote the manuscript.

